# Revealing Hidden Threats: Marine Invertebrate Invasions in the Banc d’Arguin National Park (PNBA), Mauritania, Explored Through DNA Barcoding

**DOI:** 10.1101/2024.10.18.618484

**Authors:** Carlos J. Moura, Ebaye Sidina, Ester Serrão

## Abstract

The Banc d’Arguin National Park (PNBA) in Mauritania, a globally significant biodiversity hotspot, faces growing threats from human activities and biological invasions. This study aimed to document marine invertebrate diversity in the PNBA and identify non-native and potentially invasive species. Samples were collected during expeditions in 2021 and 2022, using scuba-diving, drag dredging, and manual intertidal collection across depths of 0 to 20 meters. Through DNA barcoding of COI and 16S genes, 17 species from three phyla—Cnidaria, Bryozoa, and Arthropoda— were identified, with several cryptic taxa detected. Evidence of human-mediated introduction was found in hydroids (*Bougainvillia, Pennaria, Obelia*), bryozoans (*Amathia, Bugula, Schizoporella*), and barnacles (*Amphibalanus*), suggesting invasions from regions like Brazil, and the Indo-Pacific. Notably, *Amathia brasiliensis* and *Amathia cf. vidovici* were recorded for the first time in West Africa, alongside the detection of two other *Amathia* species, raising concerns about their potential impact on the PNBA’s biodiversity and ecosystem services. The presence of these exotic species, likely introduced via international maritime traffic and fishing activities, highlights the park’s vulnerability. The study underscores the utility of DNA barcoding for detecting cryptic diversity and tracking species dispersal, recommending urgent conservation measures and biosecurity protocols to protect the PNBA’s unique marine ecosystems.

## Introduction

The Banc d’Arguin National Park (PNBA), established in 1976, spans the western coast of Mauritania, covering 12,000 km^2^—approximately one-third of the country’s coastline. As one of the largest protected areas in West Africa, PNBA has earned international recognition as a UNESCO World Heritage Site (designated in 1989) and a Ramsar wetland of international importance (designated in 1982). The park is celebrated for its extraordinary biodiversity, which thrives in diverse habitats such as extensive seagrass meadows, mudflats, mangrove forests, islands, and desert landscapes. Together, these ecosystems form a unique land-seascape that supports a remarkable variety of terrestrial and marine species (Pottier et al., 2021; Araujo & Campredon, 2016).

The marine environment of PNBA is particularly notable for the upwelling of cold, nutrient-rich waters, which sustains one of the most productive marine ecosystems in the world. This upwelling, along with the park’s vast seagrass beds, supports a rich biological community, including several endangered elasmobranchs, commercially important fish, marine invertebrates, shorebirds, marine mammals, and sea turtles (Araujo & Campredon, 2016; Osipova et al., 2020; Chefaoui et al., 2021; Cornet et al., 2023). The presence of three key seagrass species—*Zostera noltei, Cymodocea nodosa*, and *Halodule wrightii*—further highlights the park’s ecological significance (Chefaoui et al., 2021; Trégarot et al., 2021).

Despite its ecological importance, PNBA faces growing pressures from human activities. These include overfishing, offshore oil and gas extraction, degradation of terrestrial ecosystems, and infrastructure development. Illegal fishing, particularly targeting sharks and rays, adds to the strain on conservation efforts. Additionally, climate change and pollution (such as oil spills, high concentrations of cadmium, plastic waste, and discarded fishing gear) further threaten the park’s ecosystems (Osipova et al., 2020). As a result, PNBA’s ecological integrity is increasingly under threat.

Conserving PNBA is critical not only for preserving its extremelly-rich biodiversity but also for supporting the sustainable livelihoods of local communities, particularly the Imraguen fishermen, who depend on the park’s natural resources. In this context, it is essential to document and understand the diversity of marine species, particularly invertebrates, as they serve as key indicators of the health of the park’s ecosystems.

The primary objectives of this study are to document and analyze the diversity of marine invertebrates in the Banc d’Arguin National Park, with a focus on identifying non-native and potentially invasive species. By employing DNA barcoding, this study aims to achieve precise species identification, reveal cryptic species, and clarify biogeographical distributions. For the first time, we report the presence of several exotic and invasive species in PNBA, such as cryptic hydroids and bryozoans, likely introduced through human activities. These findings emphasize the vulnerability of the park’s ecosystems to biological invasions and underscore the urgency of implementing targeted conservation measures to protect its biodiversity.

## Materials and Methods

### Sample Collection

Marine invertebrate samples were primarily collected manually by C.J.M. during a scuba-diving expedition to Banc d’Arguin National Park (PNBA), Mauritania, between November 23 and December 4, 2022, at depths ranging from 1 to 15 meters using a knife. Additionally, a bryozoan species was collected from the intertidal zone, and further specimens were retrieved using a drag dredge (“ganchorra”) during this expedition and a previous one conducted by E.S. in May 2021. Several specimens were photographed and filmed in situ to document their habitats and morphological features. Following collection, all samples were preserved in 96% ethanol, labelled with relevant metadata, and transported to the University of Algarve, Portugal, for laboratory analysis.

In the laboratory, samples were sorted by morphotype and photographed under a stereomicroscope. Tissue fragments (approximately 1–2 cm) from each distinct morphotype were isolated, re-labelled, and preserved in separate 1.5 ml Eppendorf tubes with 96% ethanol for subsequent genetic analyses.

### DNA Barcoding Analyses

The DNA barcoding protocols and bioinformatics analysis followed the methodologies outlined in Moura et al. (2024). Preliminary taxonomic identifications were made by combining morphological data with DNA barcoding results, and the taxa were cross-checked with the World Register of Introduced Marine Species (WRiMS; https://www.marinespecies.org/introduced) to identify potential non-native or invasive species (as in Moura et al., 2024).

DNA sequences for the cytochrome oxidase I (COI) and/or 16S rRNA genes were generated for the majority of species presented, except for one bryozoan. These sequences have been deposited in GenBank (accession numbers PQ374013-PQ374036, PQ374183-PQ374200). This combined morphological and molecular approach facilitated the detection of cryptic species and the identification of exotic taxa, offering valuable insights into species dispersal patterns and the risks posed by marine invasions in the PNBA ecosystem.

## Results

Seventeen marine invertebrate species, spanning three phyla—Cnidaria, Bryozoa, and Arthropoda—and associated with genera listed in the World Register of Introduced Marine Species (WRiMS) (https://www.marinespecies.org/introduced/), were recorded in the Banc d’Arguin National Park (PNBA), Mauritania (Fig. 1). These species were collected along a depth gradient from the intertidal zone down to approximately 20 meters (Table 1). With the exception of one species, all were identified through DNA barcoding, which not only confirmed their taxonomic identities but also contributed to a deeper understanding of their biogeographical distributions (Figs. S1.1-15).

**Table 1.**
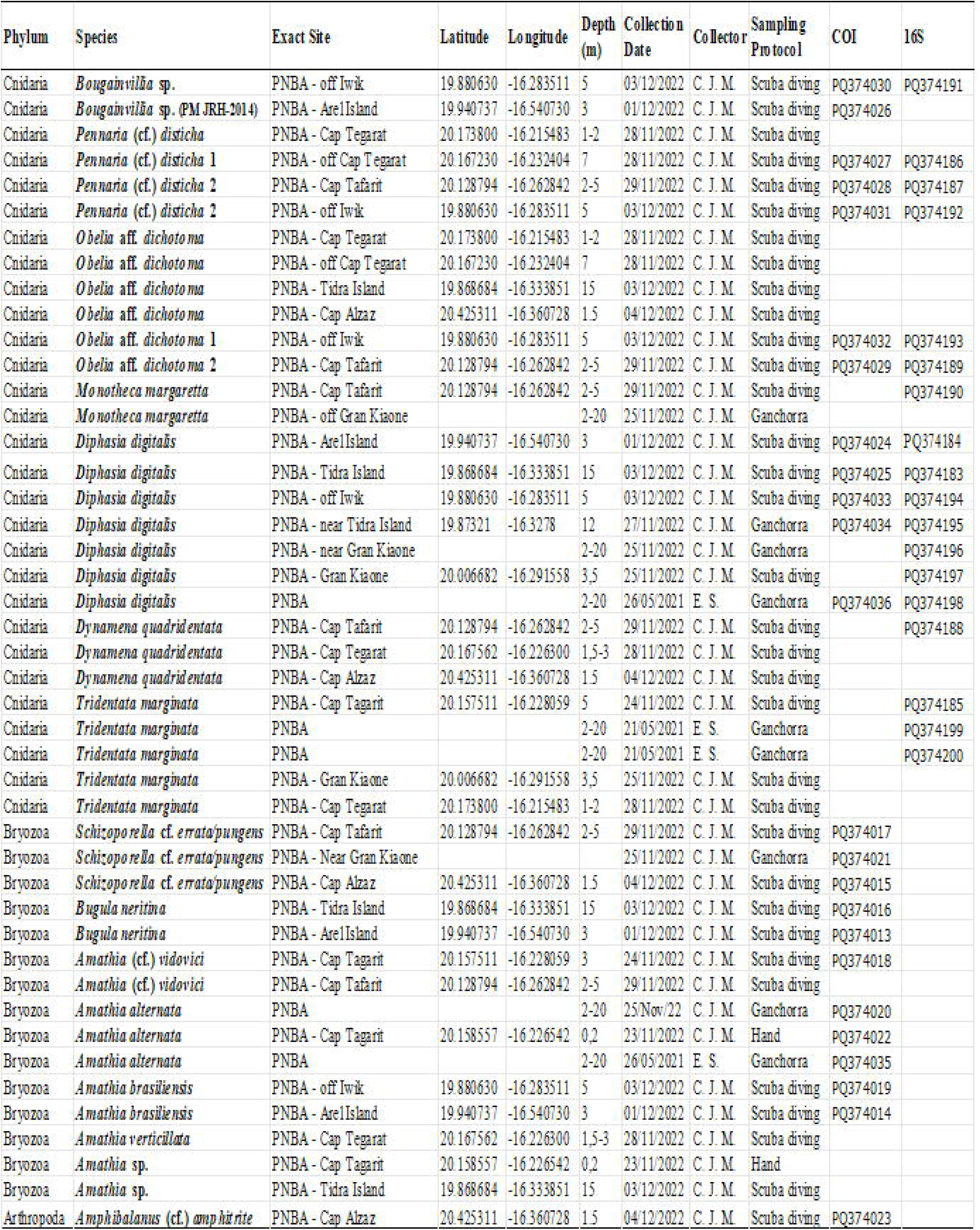
Sampling details and GenBank accession numbers of taxa identified in the PNBA.

**Fig 1.**
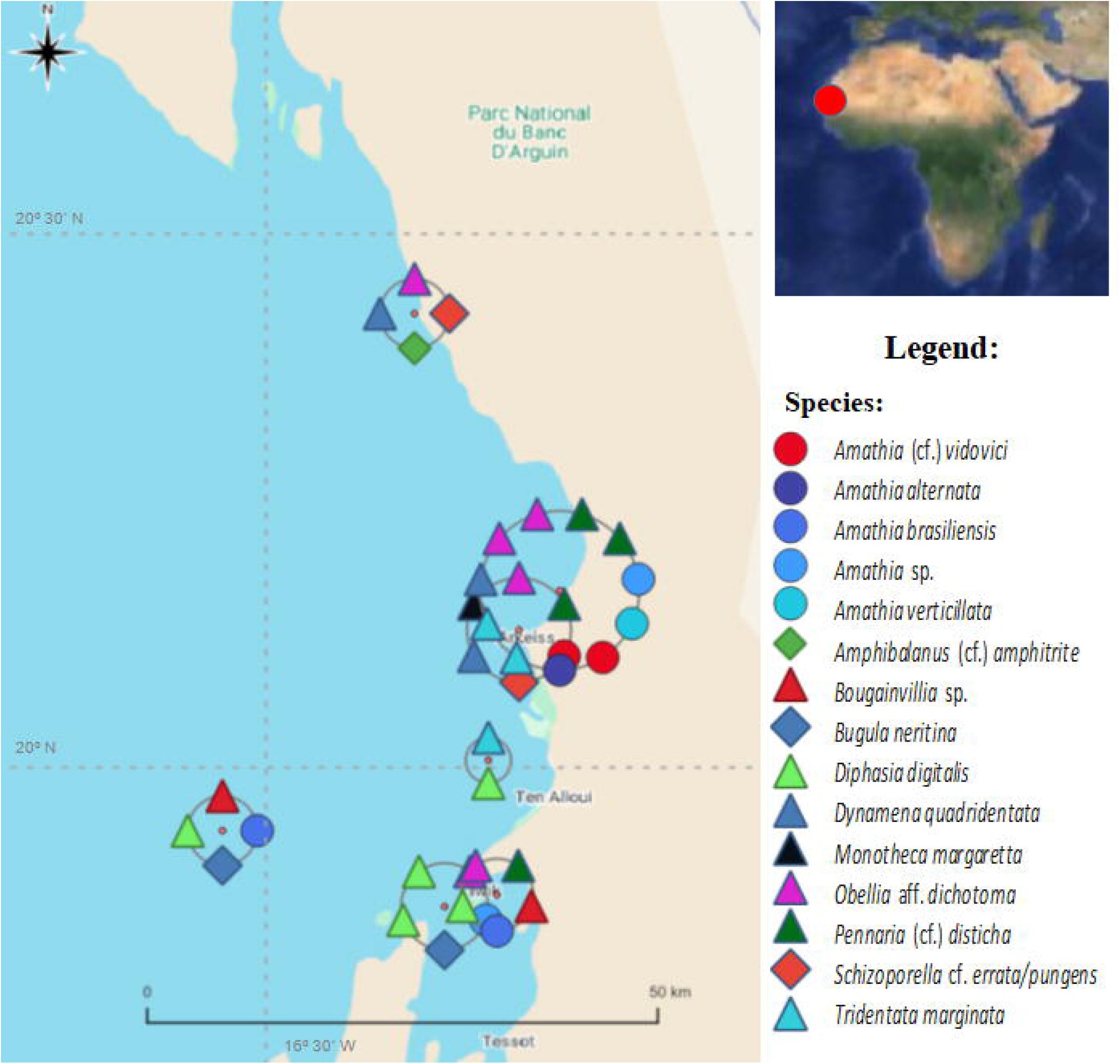
Map of sampling sites in the PNBA, Mauritania, where the identified species were found.

### Taxonomic Account and Biogeography

### Phylum Cnidaria Class Hydrozoa Order Anthoathecata

### Family Bougainvilliidae

#### *Bougainvillia* spp

(Fig. 2 A, B)

**Fig 2.**
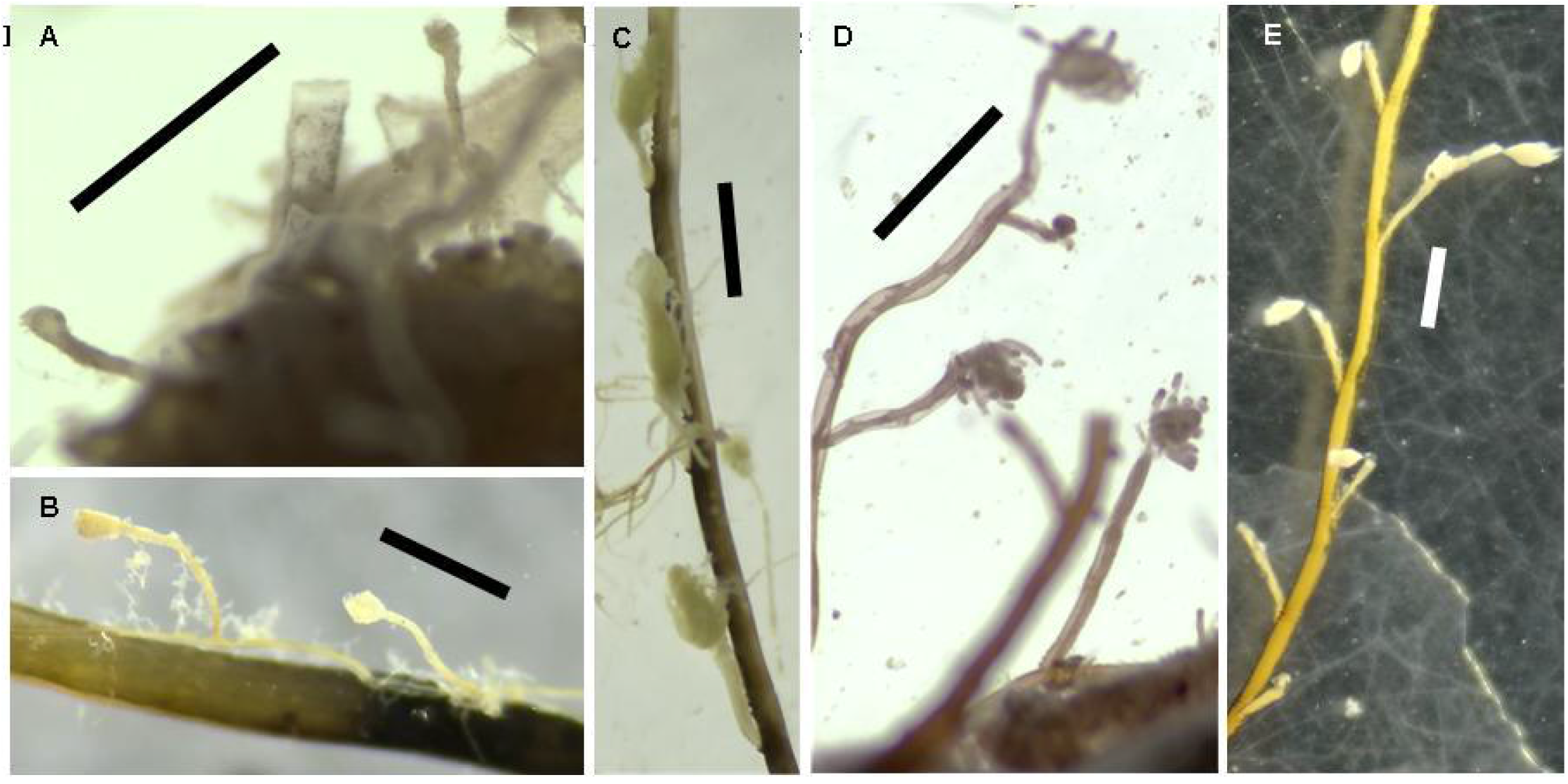
*Bougainvillia* spp. (A, B), *Pennaria* (cf.) *disticha* (C-E). Scale bars: 1mm.

Two inconspicuous species of *Bougainvillia* hydroids were sampled from the PNBA (Parc National du Banc d’Arguin): one near Arel Island, found overgrowing the hydroid *Diphasia digitalis*, and another near Iwik, overgrowing a gorgonian. DNA barcoding of the Iwik sample (Fig. 2 A) reveals it represents a distinct species of *Bougainvillia*, never previously DNA barcoded, and likely phylogenetically related to *B. muscus* (Allman, 1863) (Figs. S1.1-2). Current DNA barcode data suggest this species may fall within its natural distribution range.

In contrast, the *Bougainvillia* sample collected near Arel Island (Fig. 2 B) corresponds to a different species, previously DNA barcoded from China by He et al. (2014; GenBank direct submission: *Bougainvillia* species listed as “PM JRH-2014”; Fig. S1.1). The COI barcode of the Arel specimen differs by only one or two nucleotide positions from the Chinese specimens, indicating that this hydroid species was likely introduced to West Africa from the Indo-Pacific via human-mediated transport.

The WRiMS database (https://www.marinespecies.org/introduced/) lists four introduced species of *Bougainvillia*: *B. muscus, B. macloviana* Lesson, 1830, *B. niobe* Mayer, 1894, and *B. rugosa* Clarke, 1882. Of these, only *B. muscus* has been DNA barcoded. Phylogenetic analyses using the available 16S and COI barcodes for *B. muscus* (Figs. S1.1-2) show this species is widely distributed in the cold waters of the North Atlantic, the northeastern Pacific, and New Zealand, suggesting its spread may have been facilitated by human activity (Schuchert 2007). In addition, one cryptic species similar to *B. muscus* have been reported from warmer waters in Brazil and India (Figs. S1.1-2), likely dispersed through non-natural means too.

Based on the currently available DNA barcodes for *Bougainvillia* species (Figs. S1.1-2), we can infer marine invasions of: *B. muscus* (sensu stricto) across cold waters; a tropical cryptic species similar to *B. muscus*, found in Brazil and India, which is genetically close to the *Bougainvillia* species from Iwik, Mauritania; and an unidentified species detected both in China and near Arel Island, Mauritania.

*Bougainvillia* species have been sporadically reported in West Africa (Schuchert, 2007), including in the deep waters off Mauritania (Gil & Ramil, 2017a). However, as demonstrated by our current DNA barcoding analyses, these morphologically simple taxa are more accurately identified through genetic methods.

### Family Pennariidae

#### *Pennaria* (cf.) *disticha* Goldfuss, 1820

(Fig. 2 C-E)

The “Christmas tree hydroid” was detected four times in the PNBA: near Cap Tegarat, Cap Tafarit, and offshore from Iwik. The widely distributed nominal species *Pennaria disticha* is now recognized as a complex of at least three cryptic species (Miglietta et al., 2015; 2019; Vaga et al., 2020; Moura et al., 2024; Figs. S1.3-4)), all likely dispersed indirectly by human activity.

In Mauritania, DNA barcoding analyses (using 16S and COI markers) revealed two distinct cryptic species of *Pennaria*. The sample collected near Cap Tegarat (Fig. 2 C) belongs to a species recently identified through DNA barcoding for the first time in West Africa, specifically in the Bijagós Archipelago, Guinea-Bissau (Moura et al., 2024). This species is part of a genetic lineage also found in the Mediterranean and the Gulf of Mexico (Fig. S1.3). In contrast, the samples from Cap Tafarit (Fig. 2 D) and offshore from Iwik (Fig. 2 E) correspond to another widely distributed cryptic species of *Pennaria* (Fig. S1.3). The 16S barcodes of these two Mauritanian samples are identical to a barcode from a sample in Brazil (Fig. S1.3), suggesting a relatively recent trans-Atlantic dispersal of this invasive lineage.

*Pennaria* species are widely distributed in warm-temperate to tropical waters worldwide (Schuchert, 2006). Previous reports in West Africa include the Cape Verde Islands, Madeira, the Canary Islands, Western Sahara, Senegal, Gambia, and Guinea-Bissau (cf. Moura et al., 2024). These findings represent the first records of two *Pennaria* species in Mauritanian waters.

### Order Leptothecata

### Family Obeliidae

#### *Obelia* spp. (aff. *dichotoma* (Linnaeus, 1758)) (Figs. 3 A, B)

Specimens of *Obelia*, morphologically resembling *Obelia dichotoma*, were found near Cap Tegarat, Tidra Island, Cap Alzaz, Iwik, and Cap Tafarit. However, only the specimens from Iwik (Fig. 3 A) and Cap Tafarit (Fig. 3 B) were selected for DNA barcoding using 16S and COI markers, revealing that they correspond to two distinct species (Figs. S1.5-6).

**Fig 3.**
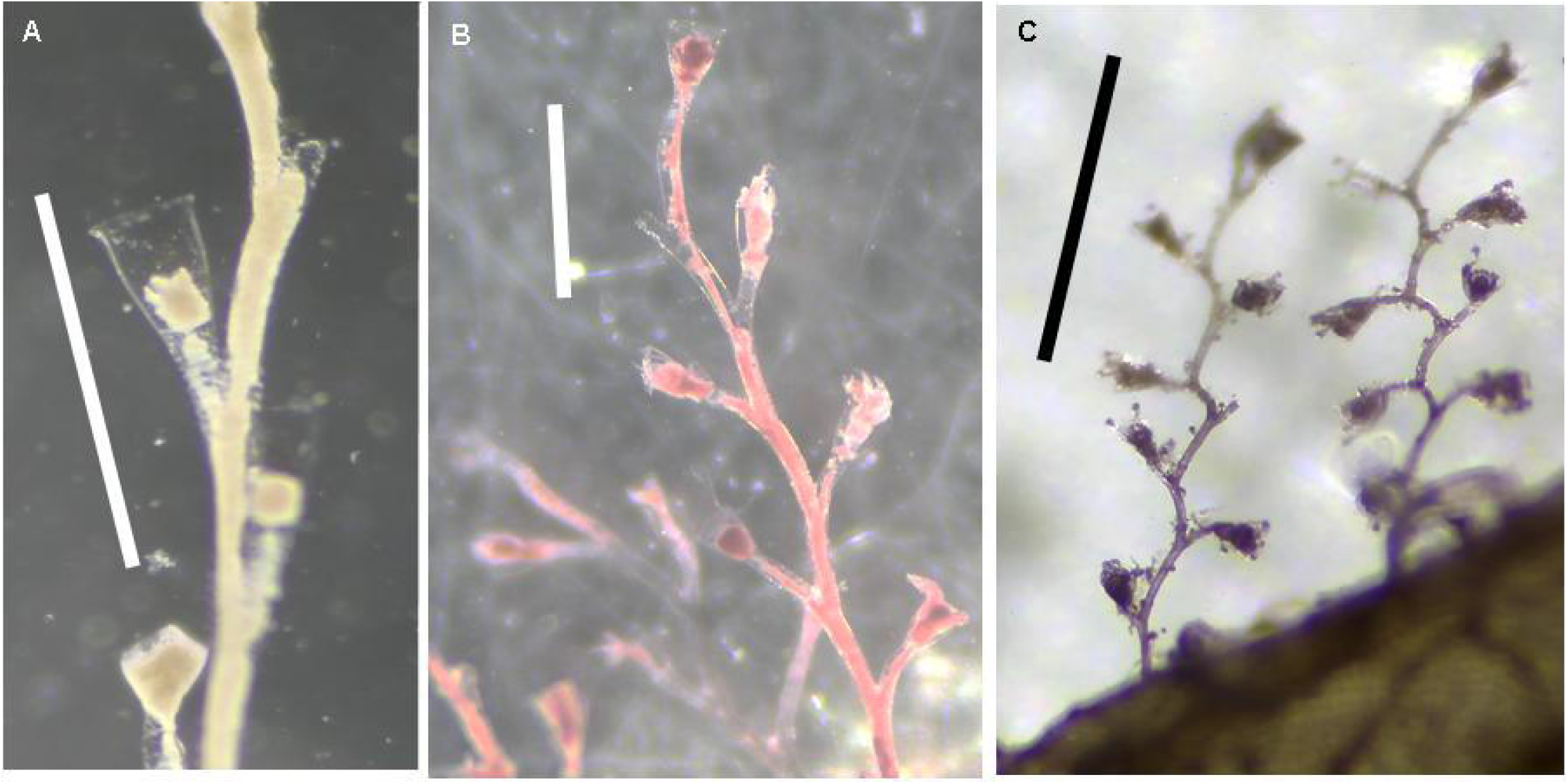
*Obelia* spp. (aff. *dichotoma*) (A, B), *Monotheca margaretta* (C). Scale bars: 1mm.

The species detected near Iwik (Fig. 3 A) belongs to a lineage previously reported in the Bijagós Archipelago, Guinea-Bissau (Moura et al., 2024) (Fig. S1.5), as well as in Hawaii (Fig. S1.6), suggesting it is an *Obelia* lineage likely dispersed through human activity.

The *Obelia* species from Cap Tafarit (Fig. 3 B), based on DNA barcoding, falls within a clade that includes specimens from Guinea-Bissau, the Gulf of Mexico (Fig. S1.5), Hawaii, and California (Fig. S1.6). The 16S phylogenetic tree (Fig. S1.5) suggests that the Mauritanian taxon may represent a distinct species, similar to the one recently identified in Guinea-Bissau (Moura et al., 2024). Further investigation into the potential human-mediated dispersal of this cryptic *Obelia* clade will require a larger set of DNA barcodes.

*Obelia dichotoma* s.l. has been frequently reported along the West African Atlantic coasts (cf. Medel and Vervoort, 2000), including in Mauritanian waters (Gil & Ramil, 2017a). However, this study provides the first evidence of two cryptic species in Mauritania with morphological similarities to *O. dichotoma*, one of which is newly detected in West Africa.

### Family Plumulariidae

#### *Monotheca margaretta* Nutting, 1900 (Fig. 3 C)

This species is reported here from near Cap Tafarit and Gran Kiaone. The 16S barcode obtained from the Cap Tafarit specimen (Fig. 3 C) indicates that it belongs to a clade of this nominal species previously identified in the Mediterranean, Azores, and Madeira (Fig. S1.7). To date, three genetic species have been identified within *Monotheca margaretta* s.l. (Moura et al., 2018). Although the available 16S barcodes for *Monotheca* are still limited, they do not yet provide conclusive evidence of human-mediated dispersal for species of this genus. Instead, the data suggest that species within *Monotheca* may be more diverse and have more restricted distributions than previously thought.

*Monotheca margaretta* s.l. has been previously reported in West Africa, including the Canary Islands, Senegal, and Ghana (cf. Ansín Agis et al., 2001). This study marks the first record of this species in Mauritania. The true *M. margaretta* appears to be restricted to the West Atlantic, whereas the *Monotheca* species found in Mauritania belongs to one of two cryptic species found in the NE Atlantic (Moura et al., 2018).

### Family Sertulariidae

#### *Diphasia digitalis* (Busk, 1852) (Fig. 4 A, B)

This species was frequently found in the PNBA, occurring at seven sampling stations. Two 16S haplotypes were identified in Mauritania, differing by a single nucleotide position, one of which was also detected in Guinea-Bissau by Moura et al. (2024) (Fig. S1.8). The COI barcodes from Mauritania were identical to those from a sample collected in Guinea-Bissau (Fig. S1.9).

**Fig 4.**
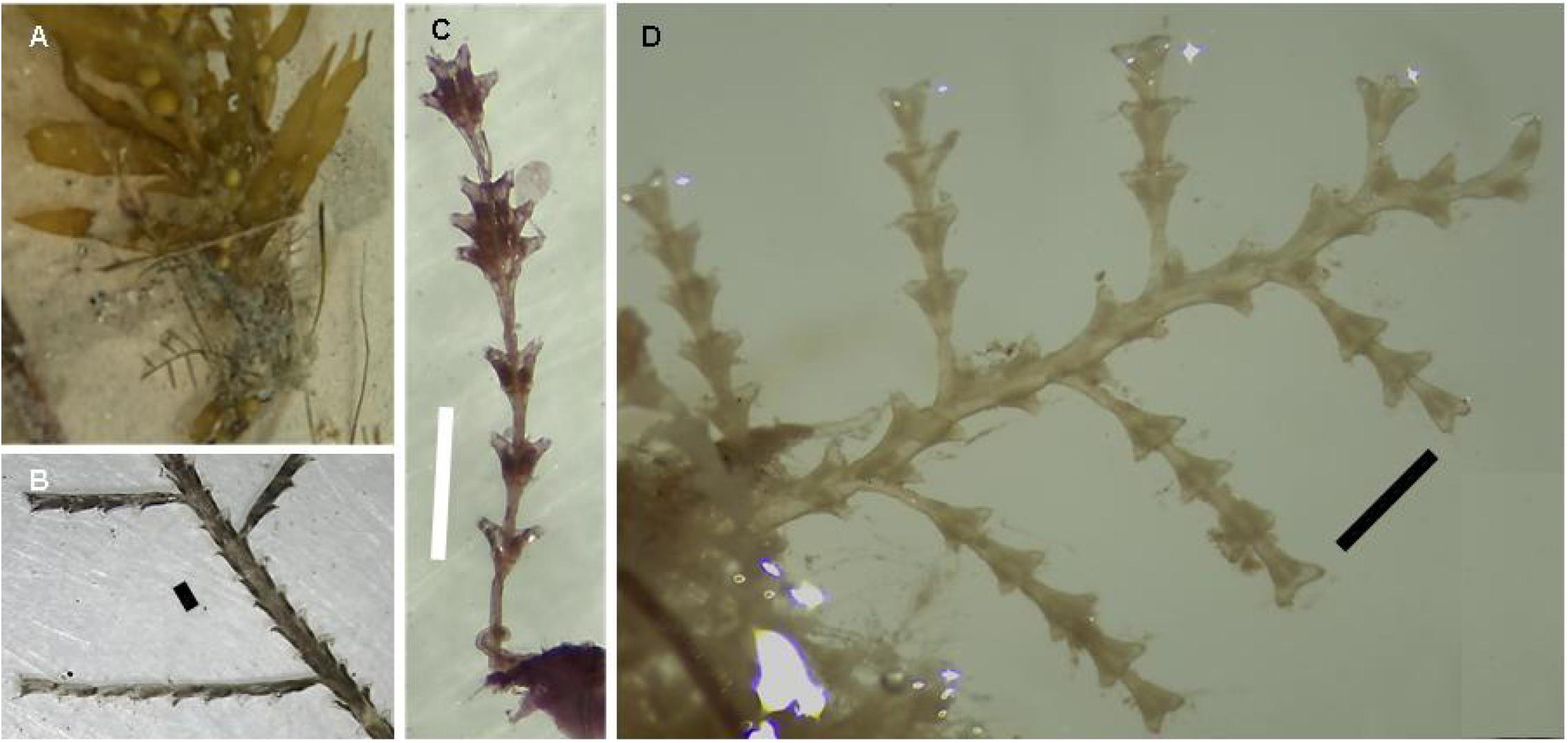
*Diphasia digitalis* (A, B), *Dynamena quadridentata* (C), *Tridentata marginata* (D). Scale bars: 1mm.

*Diphasia digitalis* is a circumglobal species, historically reported along the West African Atlantic coasts, including Guinea-Bissau, Guinea, Ivory Coast, and Gabon (Gil & Ramil, 2017b, 2023). This study marks the first record of this species in Mauritania.

#### *Dynamena quadridentata* (Ellis & Solander, 1786) (Fig. 4 C)

This taxon was discovered at three locations within the PNBA: Cap Tafarit, Cap Tegarat, and Cap Alzaz, where it was found overgrowing rocks and a sponge. The 16S barcode from one colony was identical to those of *Dynamena quadridentata* sampled in Guinea-Bissau and the Azores (Fig. S1.10), suggesting long-distance dispersal likely facilitated by rafting on algae or attachment to boat hulls (Moura et al., 2024).

*Dynamena quadridentata* has a circumglobal distribution in temperate and tropical waters (Vervoort & Watson, 2003), but has been rarely reported in West Africa (Moura et al., 2024). This study marks the first record of the species in Mauritania.

#### *Tridentata marginata* (Kirchenpauer, 1864) (Fig. 4 D)

This species was found at five sampling locations within the PNBA, primarily on rocky substrates. The 16S barcodes from three Mauritanian samples revealed two closely related haplotypes, distinct from those of samples collected in Guinea-Bissau, Madeira, the Azores, and Brazil (Fig. S1.11). Additional DNA barcodes from across its wide distribution range are needed to better understand potential biological invasions through phylogeographic analysis (Moura et al., 2024).

*Tridentata marginata* has a circumglobal distribution in tropical and subtropical seas (e.g., Medel & Vervoort, 1998) and has been frequently reported along the West African coastline, including in Mauritania (Gil & Ramil, 2017a, 2023).

### Phylum Bryozoa

### Class Gymnolaemata

### Order Cheilostomatida

#### Family Schizoporellidae

#### *Schizoporella* cf. *errata* (Waters, 1878) / *pungens* Canu & Bassler, 1928 (Fig. 5 A)

Three *Schizoporella* colonies were discovered in the PNBA at Cap Tafarit, Cap Alzaz (Fig. 5 A), and near Gran Kiaone, growing on rocky substrates. The COI barcodes from these samples correspond to the most widely distributed *Schizoporella* COI haplotype (Fig. S1.12), likely spread by human activities, and previously detected in Guinea-Bissau, the Mediterranean, and the Pacific (Moura et al., 2024).

**Fig 5.**
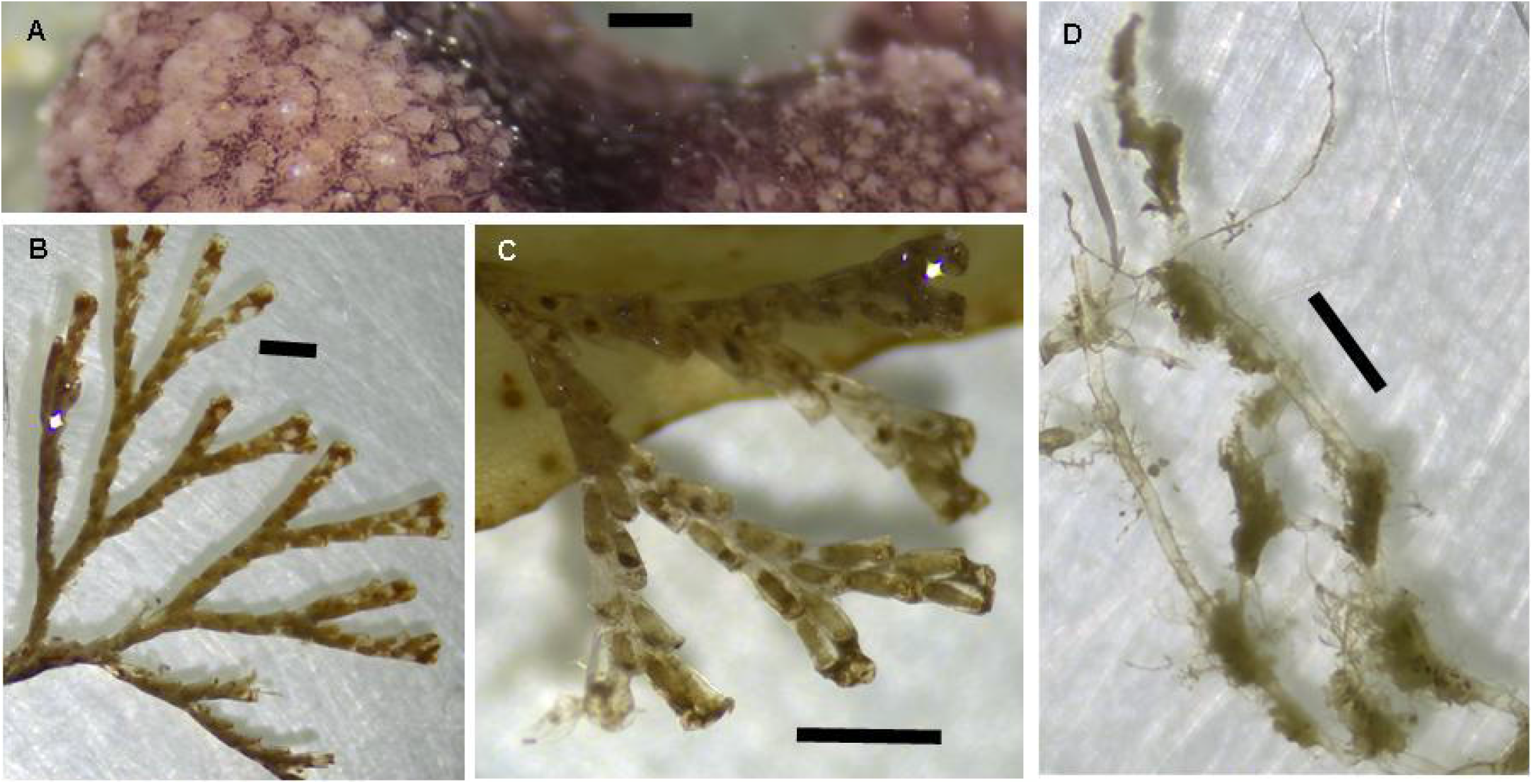
*Schizoporella pungens* (A), *Bugula* (cf.) *neritica* (B, C), *Amathia brasiliensis* (D). Scale bars: 1mm.

Both *Schizoporella errata* and *S. pungens* (sensu lato) are widespread species found in tropical and subtropical marine waters, frequently reported in ports and marinas (e.g., Marques et al., 2013; Canu & Bassler, 1930). Previous records of these two nominal species, which likely represent a single species, in West Africa (cf. Moura et al., 2024) include Angola, Ghana, the Canary Islands, Madeira, and Guinea-Bissau. This study provides the first record of this taxon in Mauritania.

### Family Bugulidae

#### *Bugula* (cf.) *neritina* (Linnaeus, 1758) (sensu lato) (Fig. 5 B, C)

The nominal species *Bugula neritina* was collected twice in the PNBA, near Tidra (Fig. 5 B) and Arel (Fig. 5 C) Islands, overgrowing a rock and an alga. The COI barcodes from these colonies correspond to the most globally widespread and common COI haplotype within the *B. neritina* species complex, which has also been detected in Guinea-Bissau, West Africa, and is known to have spread through human activities (Moura et al., 2024; Fig. S1.13).

While *Bugula neritina* s.l. has long been recorded in West Africa, including in Senegal, Madeira, the Canary Islands, Cape Verde, and Guinea-Bissau (cf. Moura et al., 2024), this study represents the first report of the species in Mauritanian waters.

### Order Ctenostomatida

### Family Vesicularioidea

### *Amathia alternata* Lamouroux, 1816

This bryozoan species was collected at three locations within the PNBA, in both intertidal and subtidal zones. The COI barcodes from these samples are identical to those from Brazil and Guinea-Bissau (Moura et al., 2024; Fig. S1.14). *Amathia alternata* had previously been documented only in the West Atlantic, from North Carolina to Brazil (cf. Nascimento et al. 2022), until this study and Moura et al. (2024) reported it for the first time in West Africa, specifically in Mauritania and Guinea-Bissau.

### *Amathia brasiliensis* Busk, 1886 (Fig. 5 D)

This *Amathia* species was detected near Arel Island (Fig. 5 D) and Iwik within the PNBA. The COI barcodes identified for these colonies are identical and closely cluster with *A. brasiliensis* lineages from Brazil and Virginia, USA (Fig. S1.14). Previously, this species was only known from the West Atlantic, specifically North Carolina, the Caribbean, and Brazil (Fehlauer-Ale et al. 2011). These two reports from Mauritania represent the first records of this bryozoan species in the Eastern Atlantic.

### *Amathia verticillata* (delle Chiaje, 1822) (Fig. 6 A, B)

The “spaghetti bryozoan” was found within the PNBA near Cap Tegarat, densely colonizing a large, abandoned fishing net. Some colonies reached over one meter in height (Fig. 6 A). Although no COI barcode was obtained for this species in this study, recent global genetic analyses show no evidence of cryptic diversity in *Amathia verticillata* (Nascimento et al. 2021). A single haplotype dominates globally, likely spread by boat traffic, while other genetic lineages are more regionally confined to the Indo-Pacific (Nascimento et al. 2021).

**Fig 6.**
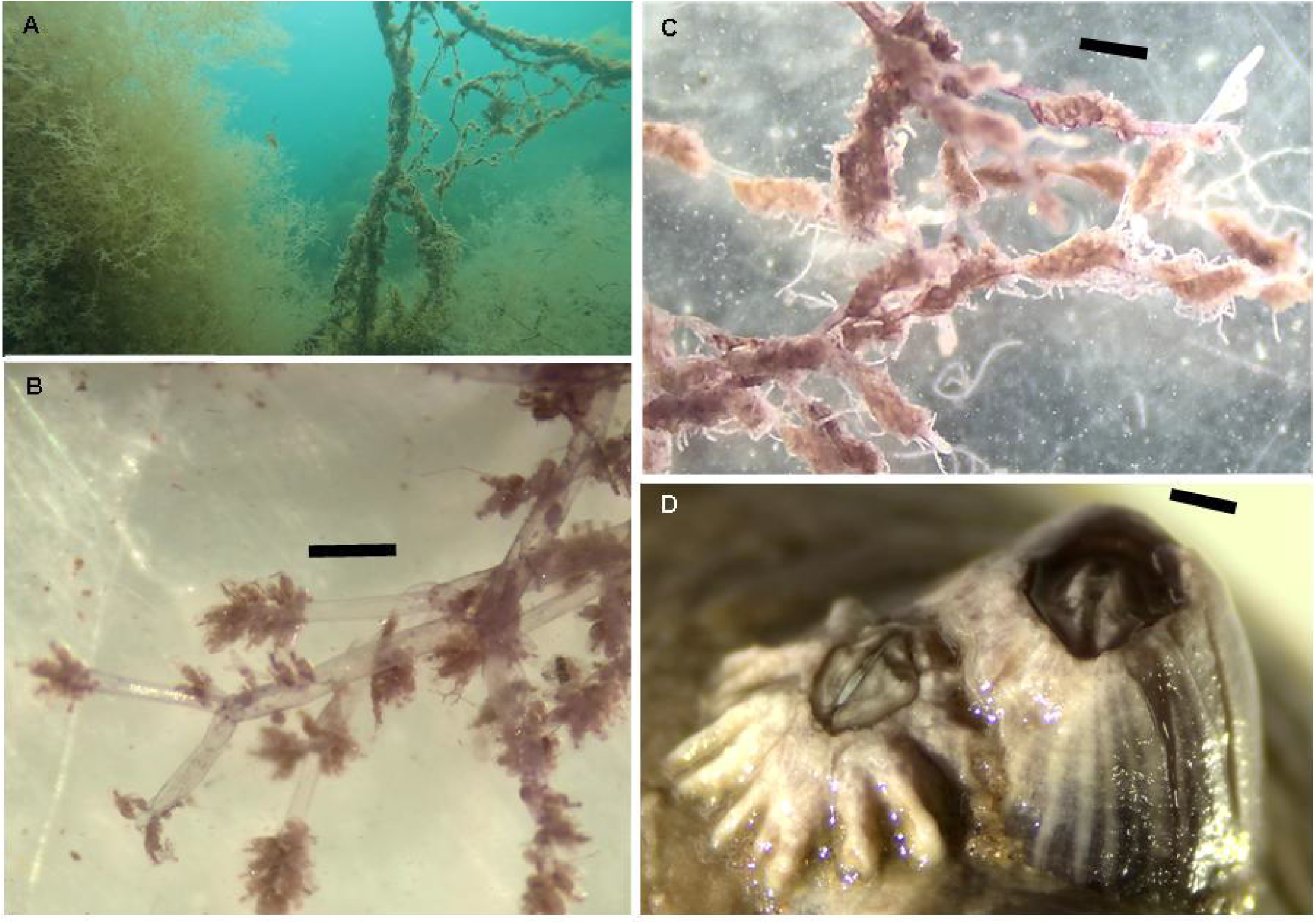
*Amathia verticillata* (A, B), *Amathia* (cf.) *vidovici* (C), *Amphibalanus* (cf.) *amphitrite* (D). Scale bars: 1mm.

Prior records of this cosmopolitan invasive species along the West African coast include Madeira, the Canary Islands, Cape Verde, Ghana, Angola, Senegal, and Guinea-Bissau (cf. Moura et al., 2024). This report represents the first documented occurrence of *A. verticillata* in Mauritania.

### *Amathia* (cf.) *vidovici* (Heller, 1867) (Fig. 6 C)

*Amathia vidovici* sensu lato was sampled twice within the PNBA, near Cap Tagarit, in the subtidal zone. This species is part of a complex (sensu Fehlauer-Ale, unpublished data, in Waeschenbach et al. 2015). Phylogenetic analysis incorporating all available COI barcodes for this nominal species reveals at least three major clades (Fig. S1.14), which likely represent different species. The COI barcode from the *A. vidovici* s.l. sample from the PNBA is identical to a barcode from Brazil, belonging to a cryptic lineage that is exclusively composed of Brazilian samples (Fig. S1.14). The shared haplotype between Brazil and Mauritania suggests a human-mediated introduction of this species.

Although *Amathia vidovici* s.l. is considered globally widespread, its true distribution remains uncertain due to cryptic diversity (Fehlauer-Ale et al. 2011). To our knowledge, this is the first report of this taxon in West Africa.

### Phylum Arthropoda

### Class Maxillopoda

### Order Balanomorpha

### Family Balanidae

#### *Amphibalanus* (cf.) *amphitrite* (Darwin, 1854) (Fig. 6 D)

A single specimen of *Amphibalanus amphitrite* s.l. was found at Cap Alzaz within the PNBA. The COI barcode from this sample clusters within the most widespread clade of this nominal species, originating from the Indo-Pacific and possessing a cosmopolitan distribution, referred to as “Clade 1” (sensu Chen et al. 2014) (Fig. S1.15). The Mauritanian sample’s COI barcode is identical to a barcode (GenBank accession: KM211413) reported by Chen et al. (2014), which could correspond to a sample from either the Indo-Pacific or the northwest Atlantic. This discovery in Mauritania indicates an expansion of *A. amphitrite* beyond its native range.

*Amphibalanus amphitrite* s.l., of Indo-Pacific origin (Carlton et al. 2015), is widely distributed globally and has been relativelly often reported in West Africa, including in Congo, Angola, Cape Verde, Sierra Leone, and the Canary Islands (GBIF 2024). This marks the first record of this taxon in Mauritania.

## Discussion

This taxonomic study of coastal marine invertebrates in the Banc d’Arguin National Park (PNBA), Mauritania, offers critical insights into the presence and spread of exotic and invasive species in the region, with considerable biogeographical and ecological implications. The research has uncovered new records of marine species and several instances of human-mediated dispersal, underscoring the park’s vulnerability to marine invasions.

DNA barcoding has confirmed that species traditionally identified through morphology, such as hydroids from the genera *Bougainvillia, Pennaria*, and *Obelia*, consist of cryptic species complexes. For example, two cryptic *Bougainvillia* species were identified: one potentially endemic and another introduced from the Indo-Pacific. Similarly, two cryptic, non-native species of *Pennaria* and two exotic cryptic species of *Obelia* were detected. Additionally, four invasive bryozoan species of the genus *Amathia* were identified, including *Amathia brasiliensis* and *A*. (cf.) *vidovici*, marking their first records in West Africa. The dispersal patterns suggest human-mediated introductions from the West Atlantic, particularly Brazil, and the Indo-Pacific regions such as China, where a cryptic hydroid species was traced back.

One of the most plausible vectors for these introductions involves international maritime traffic and fishing fleets, particularly from Asia. Fishing vessels engaged in shark-finning, which frequently operate in Mauritanian waters, likely facilitate the spread of non-native species via ballast water discharge or biofouling on hulls. Furthermore, the proximity of the PNBA to major global shipping routes and the expansion of offshore oil and gas activities in the region increases its exposure to such invasions.

These findings, along with the suggestion that marine exotic species in West Africa have been historically overlooked (Moura et al. 2024; this study), underscore the need for heightened awareness and monitoring. Neighboring coastal areas may have already acted as donor or receiver habitats for the exotic species reported here.

By contributing the first DNA barcoding evidence for various species in West Africa, this study marks an additional advancement in understanding species distribution dynamics in the region. The application of molecular tools like DNA barcoding is vital for identifying cryptic species and monitoring marine invasions, offering valuable data for biodiversity conservation and management in the face of increasing global trade and environmental change. Continued use of such methods is essential for safeguarding the unique ecosystems of West Africa.

## Acknowledgements

We thank the PNBA staff for their invaluable support during the expeditions. CJM acknowledges the field assistance of João Neiva and Mário Rolim, as well as the contributions of Ema Azevedo, Anna Santos, Emma Franke, and Filipe Nhanquê, along with other undergraduate students, for their help with sample sorting, photography, and lab work.

## Data availability

The DNA barcodes for the marine invertebrates identified from the PNBA (Mauritania, West Africa) are available in GenBank under the following accession numbers: PQ374013-PQ374036, PQ374183-PQ374200. Additional data can be provided upon request.

## Authors contributions

Expedition Participation and Organization: All authors. Sample Collection: C. J. Moura, E. Serrão. DNA Barcoding: C. J. Moura. Article Writing: C. J. Moura. Funding Acquisition: E. Serrão. Article Revision and Approval: All authors.

## Appendices

***Available at*** https://doi.org/10.6084/m9.figshare.27235734.v1

**Fig. S1.1**. Maximum-likelihood phylogenetic tree (COI marker) of *Bougainvillia* species.

**Fig. S1.2**. Maximum-likelihood phylogenetic tree (16S marker) of *Bougainvillia* species.

**Fig. S1.3**. Maximum-likelihood phylogenetic tree (16S marker) of *Pennaria* species.

**Fig. S1.4**. Maximum-likelihood phylogenetic tree (COI marker) of *Pennaria* species.

**Fig. S1.5**. Maximum-likelihood phylogenetic tree (16S marker) of *Obelia* species.

**Fig. S1.6**. Maximum-likelihood phylogenetic tree (COI marker) of *Obelia* species.

**Fig. S1.7**. Maximum-likelihood phylogenetic tree (16S marker) of *Monotheca* and *Plumularia* species.

**Fig. S1.8**. Maximum-likelihood phylogenetic tree (16S marker) of *Diphasia digitalis*.

**Fig. S1.9**. Maximum-likelihood phylogenetic tree (COI marker) of *Diphasia digitalis*.

**Fig. S1.10**. Maximum-likelihood phylogenetic tree (16S marker) of *Dynamena* species.

**Fig. S1.11**. Maximum-likelihood phylogenetic tree (16S marker) of *Tridentata marginata*.

**Fig. S1.12**. Maximum-likelihood phylogenetic tree (COI marker) of *Schizoporella* species.

**Fig. S1.13**. Maximum-likelihood phylogenetic tree (COI marker) of *Bugula neritina*.

**Fig. S1.14**. Maximum-likelihood phylogenetic tree (COI marker) of *Amathia* species.

**Fig. S1.15**. Maximum-likelihood phylogenetic tree (COI marker) of *Amphibalanus amphitrite*.

